# Genome- and Transcriptome-wide Splicing Associations with Problematic Alcohol Use and Alcohol Use Disorder

**DOI:** 10.1101/2021.03.31.437932

**Authors:** Spencer B. Huggett, Ami S. Ikeda, Qingyue Yuan, Chelsie E. Benca-Bachman, Rohan H.C. Palmer

**Author notes:** Corresponding Author: Spencer B. Huggett, Behavioral Genetics of Addiction Laboratory – Department of Psychology, Emory University, 36 Eagle Row, Atlanta, GA 30322, USA.

## Abstract

Genetic mechanisms of alternative mRNA splicing have been shown in the brain for a variety of neuropsychiatric traits, but not substance use disorders. Our study used RNA-sequencing data on alcohol use disorder (AUD) in the brain’s reward circuitry (n=56; ages 40-73; 100% ‘Caucasian’; four brain regions) and genome-wide association data on problematic alcohol use (n=435,563, ages 22-90; 100% European-American) to investigate potential genetic links with alcohol-related alternative mRNA splicing. Polygenic scores of problematic alcohol use predicted alternative mRNA brain splicing associated with AUD, which depended on brain region. Across brain regions, we found 714 differentially spliced genes in various putative addiction genes and other novel gene targets. We found 6,463 splicing quantitative trait loci (sQTLs) that were associated with the AUD differentially spliced genes. sQTLs were enriched in loose chromatin genomic regions and downstream gene targets. Additionally, the heritability of problematic alcohol use was significantly enriched for DNA variants in and around differentially spliced genes associated with AUD. Our study also performed splicing transcriptome-wide association studies (TWASs) of problematic alcohol use and other drug use traits that unveiled individual genes for follow-up and robust splicing correlations across SUDs. Finally, we show that differentially spliced genes associated showed significant overlap in primate models of chronic alcohol consumption at the gene-level in similar brain regions. Altogether, our study illuminates substantial genetic contributions of alternative mRNA splicing in relation to problematic alcohol use and AUD.

## Introduction

Alternative mRNA splicing is the process where a single gene codes for multiple mRNA transcripts and consequently multiple proteins and gene isoforms with different structures and functions. Nearly 95% of human genes undergo alternative splicing^1^. Alternative mRNA splicing in the brain is a major contributor to both the genetic and neuromolecular pathology of psychiatric traits^2^. But researchers rarely investigate genome-wide or transcriptome-wide alternative mRNA splicing associations with substance use disorders.

Alcohol consumption induces alternative splicing events^3,4^, suggesting mRNA splicing could be due to chronic alcohol use. Post-mortem human brain studies identify alternative splicing associations with alcohol use disorder (AUD) highlighting specific gene isoforms among ion channels^5^ and neurotransmitter receptors^6^ as well as intracellular pathways and synaptic plasticity processes^7^. Since individuals from these studies may be at higher genetic risk for alcohol misuse, these findings may indicate that alcohol-related alternative mRNA splicing could be due to drug exposure, genetic factors, or both.

Common genetic factors, like single nucleotide polymorphisms (SNPs; individual DNA variants), account for a modest amount of variance in problematic alcohol use^8^ and AUD^9^. While individual SNPs are associated with AUD and problematic alcohol use, these conditions are highly polygenic and share genetic risk factors with other substance use traits^10^. Outside of putative alcohol metabolism genes and neurotransmission genes, the biological basis of the genetic predisposition to AUD or problematic alcohol use remains elusive. One important mediator of genetic risk could be neuromolecular events as DNA variation has been shown to predict differentially expressed genes linked to AUD in addiction neurocircuitry^11^. How, or whether, alternative mRNA splicing mediates the genetic risk to AUD is unknown.

We hypothesized that genetic factors would 1) distinguish individuals with AUD from controls, 2) predict alternative mRNA splicing across addiction neurocircuitry, and 3) that differentially spliced genes would point to key targets underlying the genetic pathophysiology of problematic alcohol use. We also tested whether alternative mRNA splicing events were associated across brain regions and whether differentially spliced genes linked with AUD overlapped with primate models of chronic alcohol use. Our study sought to address these hypotheses and study aims using RNA-sequencing (RNA-seq) data on post-mortem human brain samples and primate alcohol use from multiple brain regions as well as large-scale genome-wide association studies (GWASs) on problematic alcohol use and other substance use traits. For an overview of our study see **Figure 1**.

**Figure 1.**
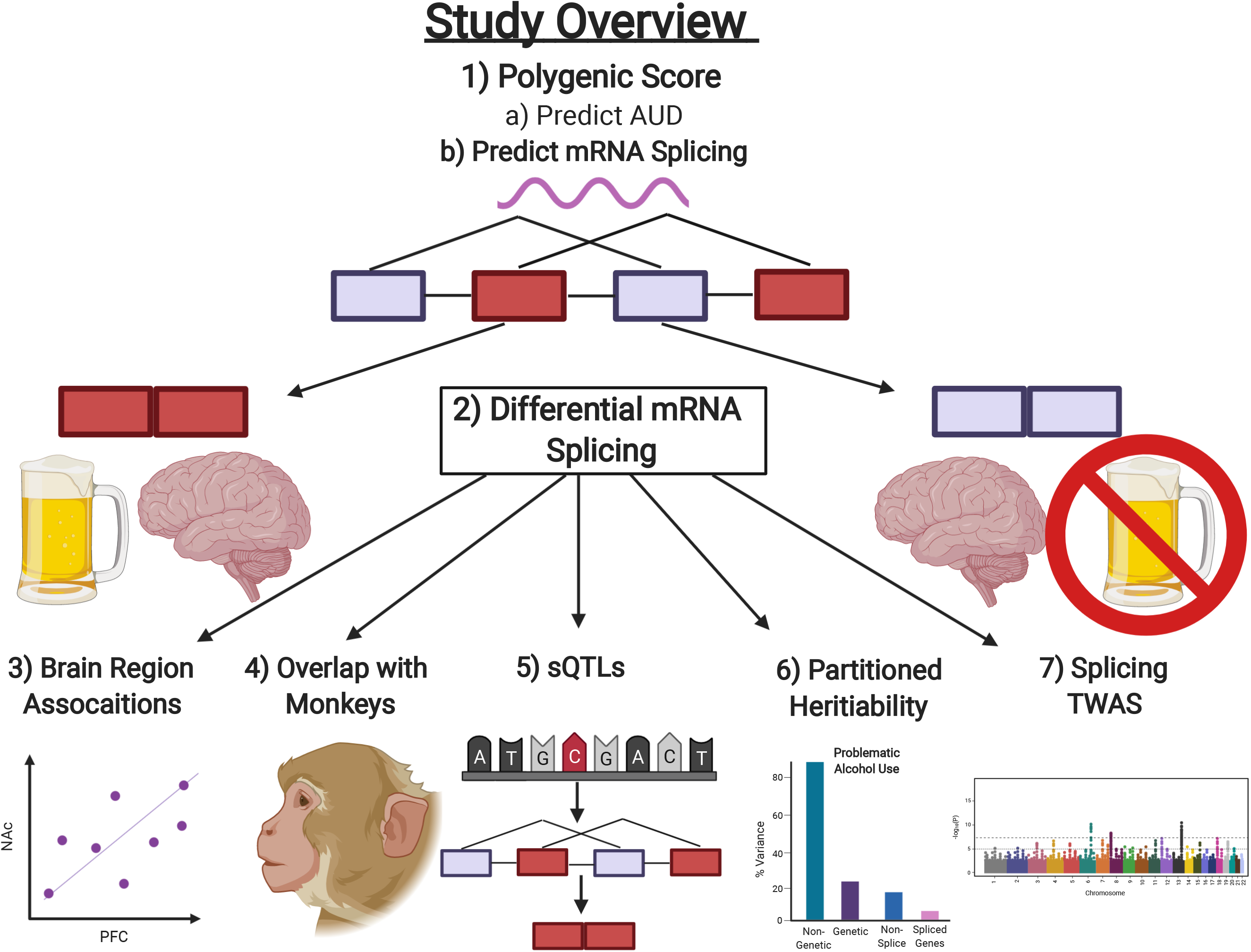
Schematic Representation of Our Study.

## Materials and Methods

### Samples

#### RNA-seq

Human post-mortem brain samples were collected from the New South Wales Brain Tissue Resource Center and included 56 genetically homogenous “Caucasian” individuals (Supplementary Figure S1). Of these samples 23.22% were female and the average age was 57.34 (s.d.=8.91, range=40-73). AUD was defined as a diagnosis of either DSM-IV alcohol abuse or dependence. Controls included social or non-drinkers that were not diagnosed with AUD and were well matched on all covariates. The most common cause of death was a cardiac complication (67.9% of all samples) followed by respiratory causes. Five individuals died of alcohol toxicity. Multiple brain regions were available for each individual and included: 1) superior pre-frontal cortex (PFC; PRJNA530758), 2) nucleus accumbens (NAc; PRJNA551775), 3) central nucleus of the amygdala (CEA; PRJNA551908) and 4) basolateral amygdala (BLA; PRJNA551909). All brain samples were collected within three days of death (post-mortem interval range=9-72 hours, *M*=32.81, s.d.=13.75 hours). For more information on this sample and RNA extraction see Rao et al. 2019^11^.

Male primate samples came from four cohorts (4, 5, 7a and 7b) of Rhesus Macaques from the Monkey Alcohol Tissue Research Resource (www.MATRR.com). Primate brain samples contained analogous brain regions as the human data, including the: 1) PFC (cortical area 32; GSE96731), 2) NAc core (GSE144783) and 3) CEA (GSE96732). Monkeys were housed individually and across cohorts had an age between 4-11 years and an average weight of 9.14 kg (s.d.=1.24). The alcohol use paradigm was described previously^12^. Briefly, monkeys were trained to drink a 4% alcohol solution for 4 months. After this, monkeys were permitted to self-administer alcohol for over a year with 22 hours of open access to alcohol. Primate alcohol consumption in this model is comparable to alcohol intake of human AUD^13^. Primate samples had five drinking categories: controls (alcohol naïve), low drinkers, high drinkers, binge drinkers or very high drinkers. To reduce multiple testing, we collapsed the top drinking categories into a single alcohol group and compared this group to the lowest drinking category (naïve controls in the NAc or the low drinking category in the PFC and CEA; note PFC and CEA samples had no naïve alcohol group). We removed samples with a normalized RNA-seq read count below two standard deviations of the group mean, which left a total of 81 primate brain samples (n_NAC_=23; n_CEA_=28; n_PFC_=30).

### GWAS

We used GWAS summary statsitics from a study on problematic alcohol use that used 435,563 individuals of European ancestry (Age range=22-90)^8^. This study collected individuals across three major cohorts: the 1) Million Veteran’s Project, 2) Psychiatric Genomics Conosoritium and 3) United Kingdom BioBank, where problematic alcohol use was defined as a DSM-V AUD diagnosis, a DSM-IV alcohol dependence diagnosis, or a log_10_ transformed metric of the Alcohol Use Disorders Identification Tests – problem drinking items.

### Data Preparation

RNA-seq data was processed using a uniform pipeline. First, we investigated RNA-seq data quality using FastQC (https://www.bioinformatics.babraham.ac.uk/projects/fastqc/). We removed Illumina adapters and poor quality reads (reads<36bp long, leading or trailing reads<Phred score of 3 and allowing a maximum of 2 mismatches per read) using Trimmomatic (version 0.39)^14^. Then, we aligned trimmed reads to either the human hg19 genome or the Rhesus Macaque genome (mm10) using STAR aligner version 2.5.3.a^15^. We followed the guidelines outlined by leafcutter (https://davidaknowles.github.io/leafcutter) to align RNA-seq reads and prepare data for differential splicing analyses. RNA-seq read alignment yielded an average of 78,955,738 reads in humans (s.d.=29,804,777; *M*_Alignment_=86.16%) and a mean of 34,551,920 reads in primates (s.d.=8,202,258; *M*_Alignment_=79.71%).

Human RNA-seq data were phased and imputed with Beagle version 5.1, which uses a probabilistic Hidden Markov Chain model that performs well for sequencing data with sparse genomic coverage^16^. Our analyses used standard methods for quality control including: genotyping rate > 95%, minor allele frequency > 0.10, Hardy-Weinberg equilibrium < 1e-6, read depth > 5 reads per sample, Phred Score > 20 and an imputation score > 0.3. After imputing the AUD data to the 1K Genomes Phase III all data, there were between 137,073-158,856 SNPs for sQTL analyses (# of SNPs depended on brain region).

We applied standard quality control to the problematic alcohol use GWAS summary statistics selecting biallelic DNA variants with a minor allele frequency > 1% and imputation information score > 0.80 while also removing ambiguous and duplicate SNPs^17^ (as recommended by: https://choishingwan.github.io/PRS-Tutorial/base/).

### Analyses

#### Differential Splicing

To detect alternative mRNA associations with AUD we used Leafcutter version 0.2.9^18^. Leafcutter is a powerful transcriptome-wide splicing method that uses a Dirichlet-multinomial generalized linear regression to identify differentially spliced genes. A differentially spliced gene is composed of multiple clusters, each of which includes a number of alternative splicing events, such as: exon skipping (see **Figure 1**), intron retention, alternative acceptor or alternative donor splice sites, which we annotated with the Vertebrate Alternative Splicing and Transcription Database (https://vastdb.crg.eu/wiki/Main_Page). Each splicing event corresponds to a change in percent spliced in (ΔPSI or dPSI) metric. In our AUD analyses, a positive ΔPSI for an exon skipping event would suggest that an individual with AUD is more likely to skip a certain exon than someone without AUD. We utilized the default filtering parameters of Leafcutter that filtered out splicing clusters with < 5 samplers per intron, < 3 samples per group and required at least 20 reads, which resulted in 18,685 unique genes across human brain regions. Differentially spliced genes/clusters were those that survived a standard Benjamini-Hochberg False Discovery (BH-FDR) rate < 0.05. Since only 21 genes were differentially spliced in primates (BH-FDR<.05), we defined significant differential splicing with a nominal p-value threshold < 0.05. We assessed linear correlations of the ΔPSI across all significant alternative splicing events that were common across brain regions. To assess the overlap between human and primate results we used a Fisher’s Exact test at the gene-level and restricted analyses to homologous genes identified by biomaRt^19^ and only used results from analogous regions of the brain (CEA, NAc and PFC). In humans, we compared our differential splicing analyses with differentially expressed genes using DESeq2 software^20^ and the same covariates and p-value adjustment.

#### Polygenic scores

We investigated two questions with polygenic score analyses. First, were human brain samples with AUD at higher genetic risk for problematic alcohol use? Second, does genetic risk for problematic alcohol use predict alternative mRNA splicing in the brain? Our study used PRScice.2 (version 2.3.3)^21^ and elected to use standard polygenic score guidelines ^17^, but decided not to use clumping given sparse genotypic data from RNA-sequencing. As a sanity check, we re-ran polygenic analyses with clumping and found similar results (see Supplementary Figure S2).

#### sQTLs

A splicing quantitative trait locus (sQTL) is a SNP that predicts alternative mRNA splicing associated with a trait. Similar to Li et al.^18^, we standardized excision-splicing ratios and then quantile normalized splicing data across individuals. Our analyses used default settings on MatrixQTL to find cis acting sQTLs that may affect mRNA splicing in a nearby gene, which tests all SNPS within 1 megabase (Mb) of a genomic region. sQTLs were defined as a SNP associated with a differentially spliced gene that survived a BH-FDR correction for multiple testing per SNP. To determine whether sQTLs resided in specific regions of the genome we annotated sQTLs in 11 annotation categories from ANNOVAR (version 4.1)^22^. The annotation categories that were built on hg18 genome coordinates were updated to their corresponding hg19 values using CrossMap (version 0.5.1)^23^.

#### Partioned Heritability

To test whether differentially spliced genes associated with AUD in the brain pointed to genetic mechanisms of alcohol misuse we performed a partitioned heritability analysis. We used LD score regression and created an annotated gene-set of differentially spliced genes (BH-FDR<.05). To be consistent with our sQTL analyses, this included SNPs within 1 Mb of the start and stop site of a differentially spliced gene, which is similar to defaults on other splicing partitioned heritability mapping tools (e.g., Li et al.^24^). To determine the specificity of our findings, we tested the partitioned heritability of this gene-set with a negative control trait (Joint disorders found via: http://www.nealelab.is/uk-biobank) that used individuals of European ancestry and had similar sample size (n=~361,194), and trait heritability (*h*^2^_SNP_=0.0695) as problematic alcohol use.

### Splicing TWASs

We performed transcriptome-wide association studies (TWASs) via splicing SMulti-Xcan^25,26^, to assess how DNA associations predicted alternative mRNA splicing associations in human tissues. To increase power, we selected all 49 Genotype Tissue Expression (GTEx) database tissues (which included up to 838 human donors; https://www.gtexportal.org/home/) as done previously^2^. We also report results from our splicing TWAS on problematic alcohol use incorporating only the 13 GTEx brain tissues, which yielded similar results (see Supplementary File S1). SMulti-Xcan combines multiple regression and elastic neural networks to predict alternative mRNA splicing from cis-sQTLs. This method accounts for linkage disequilibrium (LD) of European ancestry using the 1K Genomes Phase 3 data. Our study assessed the convergence between the splicing TWAS on problematic alcohol use and the differentially spliced genes in the brain associated with AUD. Of the overlapping genes, we assessed SNP associations mapped to these genes that were associated with other traits via https://www.ebi.ac.uk/gwas/. For these genes that also had a significant sQTL we evaluated the LD between the lead sQTL SNP (smallest p-value for the gene) with the SNP listed in the GWAS catalogue using LDlink (European Ancestry; https://ldlink.nci.nih.gov/?tab=home). Lastly, we investigated how splicing associations generalized across substance use traits by correlating splicing TWAS results from three other GWASs: cigarettes per day (n=263,954)^27^, opioid use disorder (n=82,707)^28^ and cannabis use disorder (n=374,287)^29^.

## Results

### Polygenic scores

Polygenic score analyses indicated that individuals with AUD were at higher genetic risk for problematic alcohol use than those without AUD (p=0.030; **Figure 2A**). Polygenic scores were predictive of alternative mRNA splicing but this depended on brain region (**Figure 2B**).

**Figure 2.**
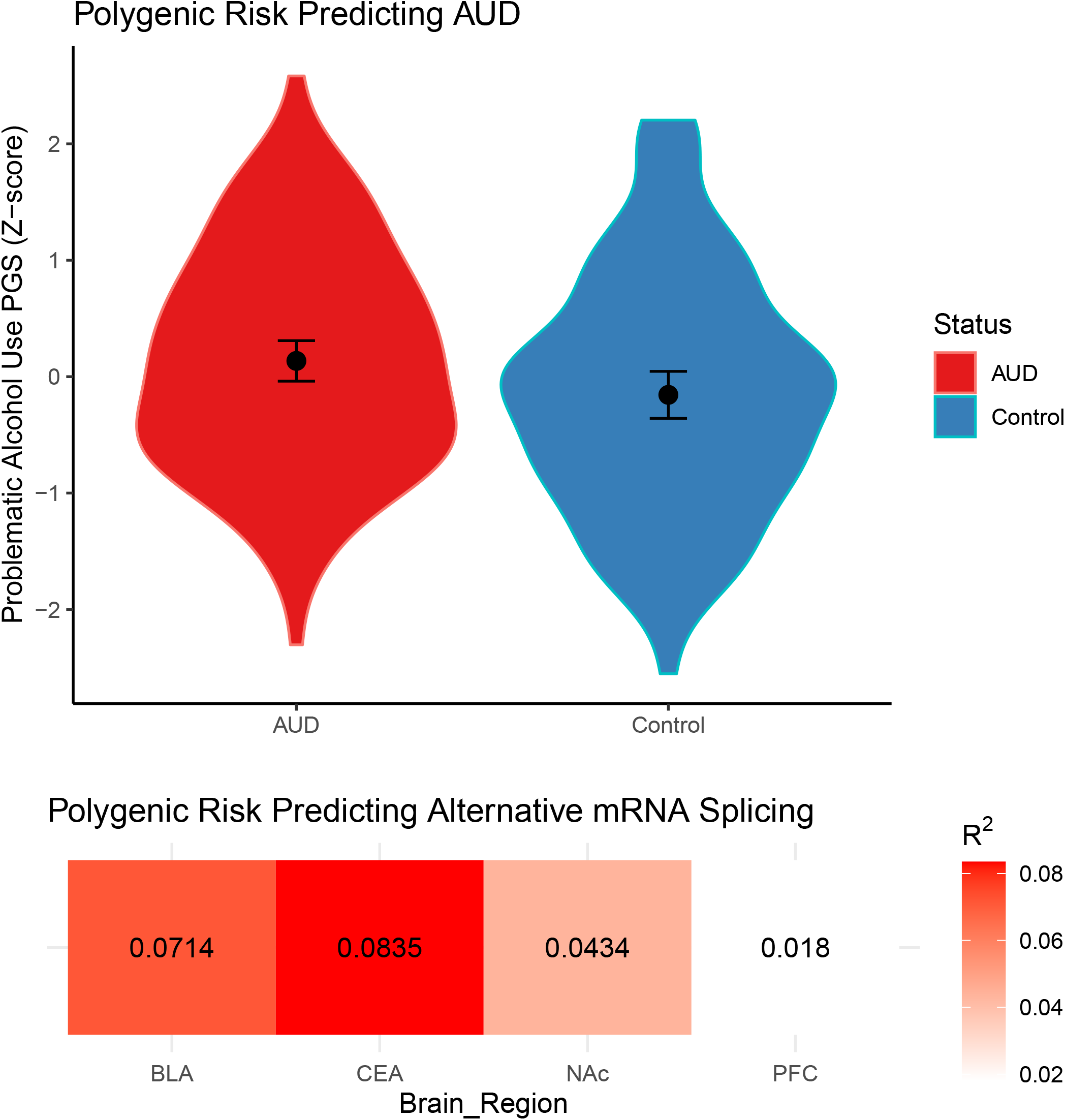
Genetic Risk for Problematic Alcohol Use and Alternative mRNA Splicing. **A)** Violin plot showing polygenic score distributions of problematic alcohol use between individuals with AUD and controls. Mean and standard error are shown. **B)** Heat matrix showing the amount of variance explained (R^2^) by polygenic prediction of differential splicing results for each brain region. Principal component (PC) analysis was used to distil differential splicing results into a single metric (1^st^ PC) that explained less than 5% of the variance in AUD-related splicing for each brain region (1^st^ PC_BLA_ = 4.364%; 1^st^ PC_CEA_ = 2.922%; 1^st^ PC_NAc_ = 2.32%; 1^st^ PC_PFC_ = 2.951). For results of polygenic score analyses with clumping see Supplementary Figure S2.

### Differential Splicing

In total, we found 714 differentially spliced genes in 740 clusters encompassing 5,118 unique splicing events associated with AUD (see **Figure 3A** & Supplementary File S2). Similar to previous analyses with these data, 92.3% of the reported differentially spliced genes associated with AUD^7^, were at least nominally significant in our analyses. From our differential splicing analyses, we identified exon skipping as the most frequent splicing event (53.9%) and found alternative splice donor events (4.0%) to be the least frequent. Differentially spliced genes were not enriched for gene ontological processes (all *p*_adj_>0.39), but several addiction genes were found to be differentially spliced, including: *ALDH3A2, CAMK2D, CAMKK2, GRIA2*, *GRK4, GRK6, HDAC3, PPP2R1B* and *PRKACB* (see Supplementary Figures S3-S4). The *GRIA2* gene showed differential splicing in a putative ‘flip flop’ splicing site (see Supplementary Figure S5), which alters the rate of AMPA receptor opening^30, 31^ and has been implicated with chronic alcohol use in primates^32^. Notably, we found no differentially expressed genes associated with AUD for any brain region (all p>0.0012, all p_adj_>0.999; see Supplementary File S3).

**Figure 3.**
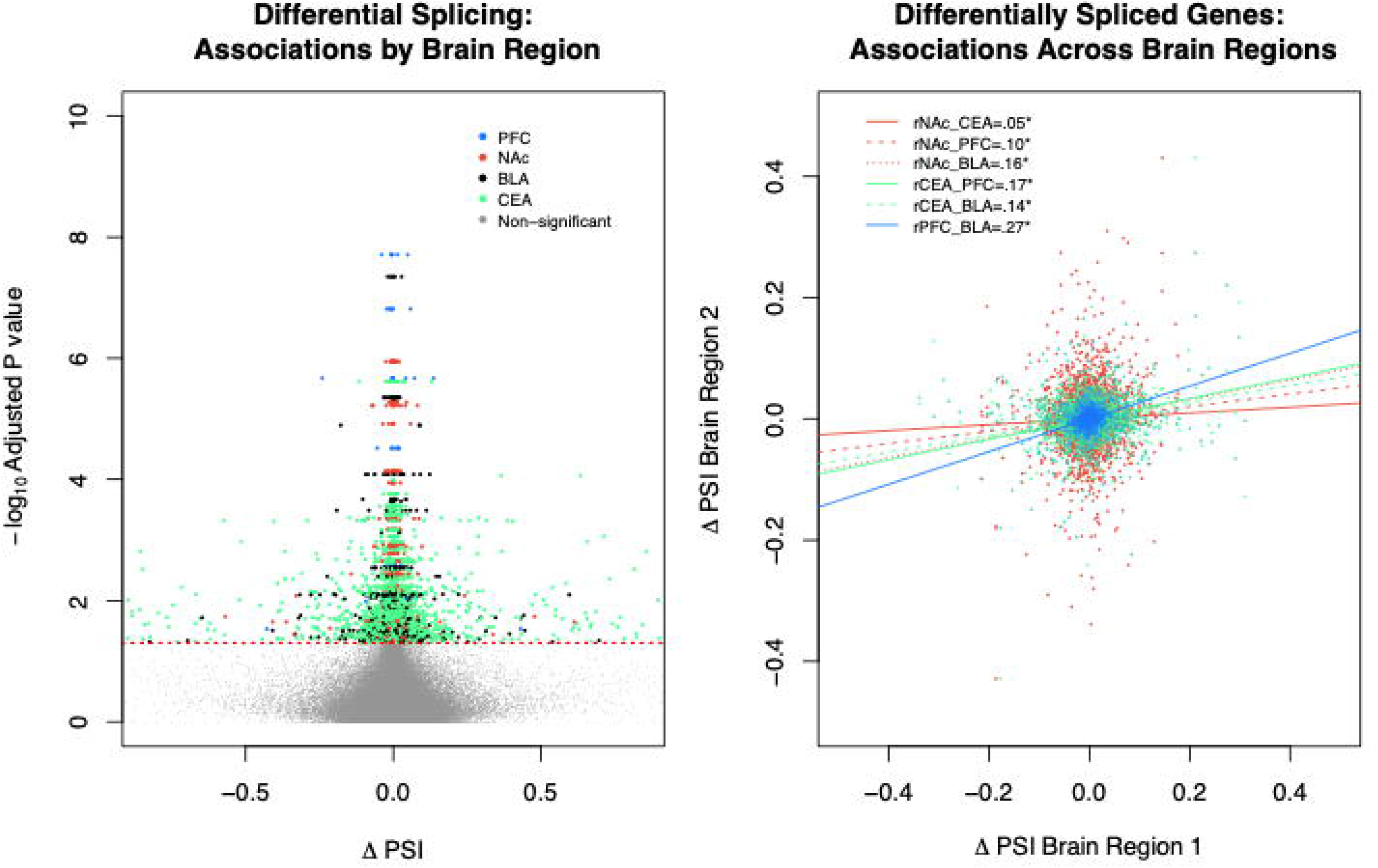
Alternative mRNA Splicing Associations with AUD by Brain Region. **A)** Volcano plot displaying differentially spliced genes between individuals with AUD and controls for each brain region. **B)** Scatter plot showing differential splicing associations across brain regions from differentially spliced genes. Note ΔPSI stands for the change in percent-spliced-in and that each colored dot represents a specific splicing event in a cluster from a significantly differentially spliced gene (*p*_adj_ < 0.05)

Investigating analogous brain regions in Macaques, we found that AUD differentially *spliced* genes tended to also demonstrate differential splicing in primate models of chronic alcohol use (see Supplementary Figure S6). This overlap was more than we expected by chance, OR=1.38, 95% CI [1.06, 1.77], *p*=0.0126. We found significant, yet small, correlations of splicing events across brain regions in humans (r=0.05–0.27; see **Figure 3B**) with the largest associations observed with the BLA. In the primate data, which lacks the BLA, we found splicing event associations between PFC and CEA (*r*=0.10, *p*=2e-16), but negative associations between the NAc with the PFC (r=-0.04) and NAc with CEA (r=-0.08, all p<0.002).

### sQTLs

Next, we tested for sQTLs, or whether specific genetic variants predicted the differentially spliced genes associated with AUD. In total, we found 6,463 unique sQTLs associated with 170 different genes (p_adj_<0.05; see **Figure 4A** and Supplementary File S4). Drug metabolism (*CYP2C19* and *CYP2C9*) intracellular signaling (*GRK4, GRK6, HDAC3, PRKACB* and *MAPK3K6*) and calcium ion channel genes (*CACNA1A, CACNA1G, CACNB2* and *KCNMA1*) had sQTL(s). Exon skipping events in the *CACNA1A* and *KCNMA1* genes corresponded to certain gene formations that differentially alter vesicular release^33^ and activation of Ca^+^ channels^34^. Most sQTLs were located in intergenic regions (52.3%) or introns (36.1%), but we only identified sQTL enrichment among DNaseI hypersensitivity sites and downstream locations of protein coding genes (see **Figure 4B**).

**Figure 4.**
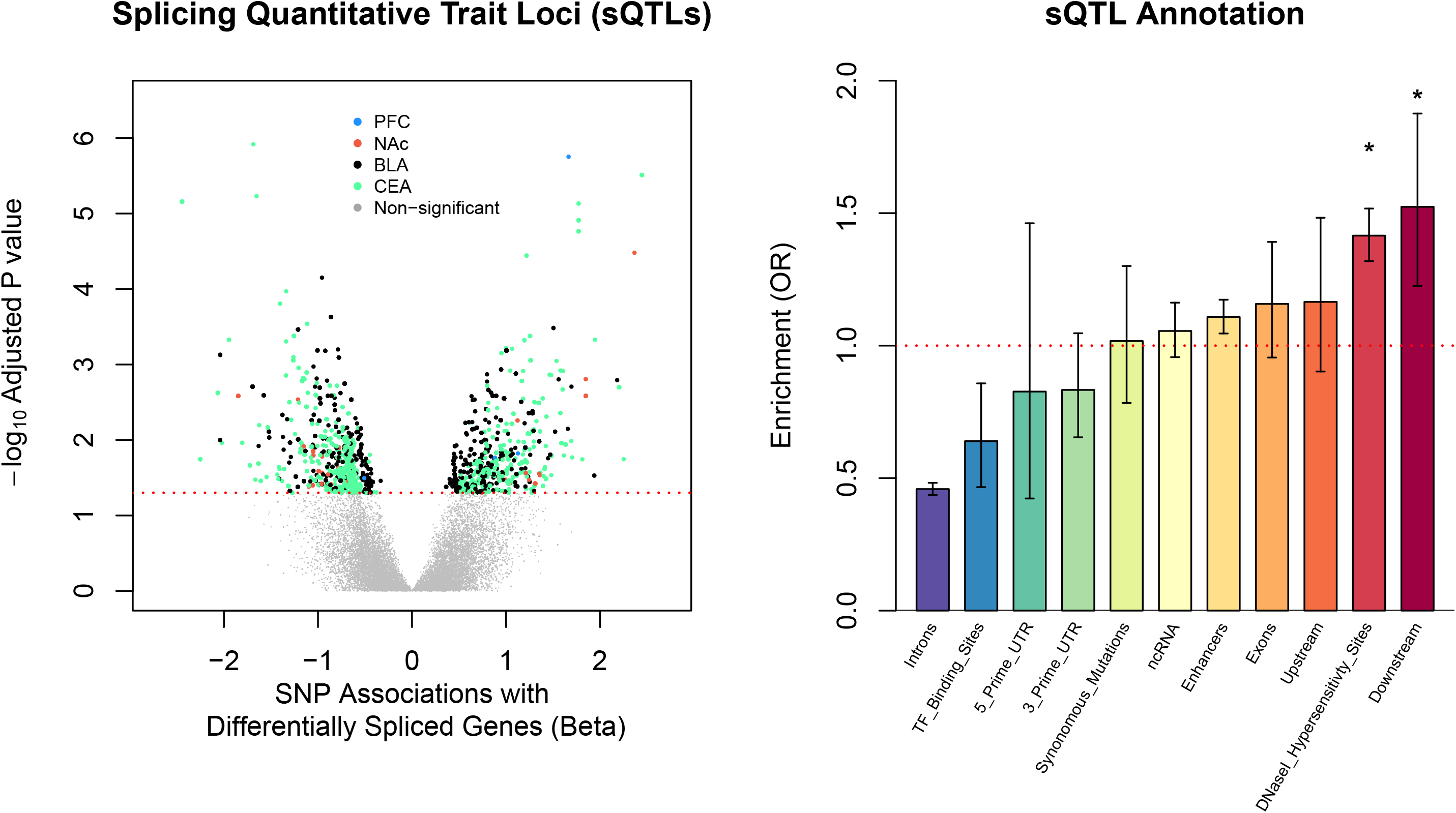
Individual DNA Markers Predict Alternative mRNA Splicing Events Associated with AUD. **A)** Volcano plot showing results from our sQTL analyses. Each dot above the dashed red line represents a significant (*p*_adj_ < 0.05) SNP association with a differentially spliced gene. **B**) Bar plot showing the genomic regions enriched for significant sQTL associations. * indicated that a certain genomic region survived correction for multiple testing (*p*_adj_ < 0.05).

### Partitioned Heritability

We further investigated the role of alternative splicing for the genetic basis of problematic alcohol use. Using LDscore regression we observed that heritable influences explained 7.81% of the individual differences in problematic alcohol use. Our partitioned heritability analyses revealed that SNPs in and around differentially spliced genes accounted for 30% of the genetic risk for problematic alcohol use (OR=1.349, se=0.064, p=6.46e-7; see **Figure 5**), but not for our negative control trait (Joint disorders, *p*=0.161).

**Figure 5.**
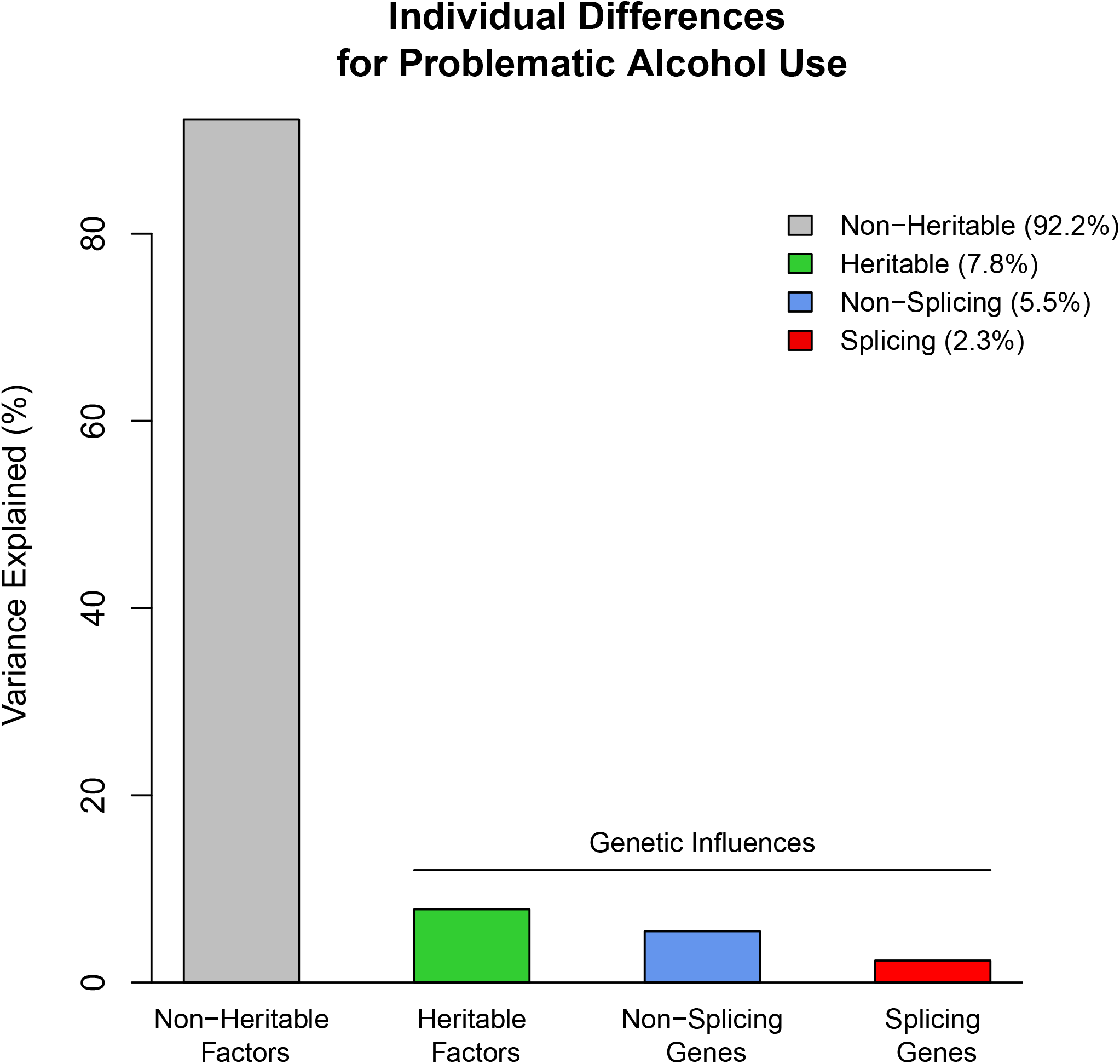
SNPs Within and Around Differentially Spliced Genes Contribute to the Heritability of Problematic Alcohol Use. Heritable factors include the observed heritability from LD score regression analyses. Splicing genes include all biallelic SNPs within and 1 Mb around the transcription start and end site of differentially spliced genes associated with AUD in the brain.

### Splicing TWASs

We found 311 splicing TWAS associations with problematic alcohol use (p_adj_<0.05; 215 unique genes; see Supplementary File S5), which were enriched for alcohol dehydrogenase activity (p_adj_=3.23e-10). Seven of the TWAS splicing genes were also differentially spliced genes in post-mortem brain tissue (*GRK4, KLHDC8B, PDS5A, PSMD7, TMEM184B, VRK2* and *WDR27*). The role of these genes in the pathophysiology of AUD is largely unknown. Previous research suggests that SNPs mapped to these genes are associated with substance use traits, neuropsychiatric illnesses and neurological endophenotypes as well as other unrelated traits (see Supplementary File S6). Of note, our lead sQTLs for the *GRK4* (rs2858038) and *KLHDC8B* (rs3819325) genes were in LD with SNPs associated with human cigarettes per day (rs2960306, R^2^=0.29) and smoking cessation (rs7617480, R^2^=0.07)^27^. To investigate potential shared genetic processes across substance use, we correlated significant splicing TWAS associations across three substance use traits: cigarettes per day^27^, opioid use disorder^28^ and cannabis use disorder^29^. Using the 1,397 significant splicing TWAS associations across substance use traits (BH-FDR<0.05; 923 unique genes; see Supplementary File S7), we found substantial overlap – especially among disordered substance use (all r>0.38; see Supplementary Figure S7).

## Discussion

Our study found novel genome-wide and transcriptome-wide splicing associations with problematic alcohol use and AUD. We found support for our three hypotheses, such that: genetic factors predicted 1) AUD, 2) alternative mRNA splicing in the brain and 3) that DNA variants in and around differentially spliced genes contributed to the heritability of problematic alcohol use. Altogether, we used a handful of methods that provided consistent evidence implicating genetic factors in AUD-related alternative mRNA splicing. These data add another layer to the neuroepigenetic understanding of compulsive alcohol use.

Extending research on other neurological traits^24,34^, we show that individual genetic markers (sQTLs) and polygenic risk underlie alternative mRNA splicing associated with AUD. Similar to other research^35^, we found that sQTLs were enriched among DNaseI hypersensitivity sites, corroborating that lose chromatin regions are hotspots for alternative mRNA splicing regulation. Previous splicing studies used a single tissue type^2,35–37^ and our study extends this work and encourages future work to investigate multiple tissue types when possible - as the genetic links with splicing events may differ by brain region.

Splicing associations with alcohol misuse occurred in genes involved with neurotransmission, intracellular signaling and drug/alcohol metabolism. Most alternative mRNA splicing events were uncharacterized, but a few of the ion channel (*CACNA1A, KCNMA1*) and glutamate receptor (*GRIA2*) associations seemed to affect synaptic neurotransmission. For instance, in the BLA, we found that individuals with AUD were more likely to have an exon skipping event of the *GRIA2* flip exon (exon 14), which is associated with longer glutamate receptor opening and consistent with the BLA pathology in alcohol use^38–40^.

The takeaways from our study are in accordance with previous analyses with these data. While we found a different number of significant AUD associations in human brain tissue – likely due to methodological differences – we found that differential splicing had more significant associations with AUD than differential expression^7^ and that splicing associations most frequently incurred exon skipping events and were largely tissue specific. Also, at the genetic level, we identified an order of magnitude more sQTLs than the previously reported (and validated) expression QTLs (eQTLs) with AUD^11^. These results are in accordance with previous analyses suggesting alternative mRNA splicing elicits robust genetic and neurotranscriptional correlates with psychiatric traits^2^ and calls for additional research to better characterize the gene isoform architecture of mental illness and substance abuse.

We found *preliminary* evidence that alternative mRNA splicing could play a more general role in a common genetic liability of substance use disorders and psychopathology. Our study revealed moderate splicing associations across disordered and problematic drug use as well as tobacco consumption, via splicing TWASs. Furthermore, the sQTLs underlying AUD-related differential splicing in the brain were correlated with DNA variants previously implicated in tobacco consumption, mental illness, and cognitive functioning. Additionally, differentially spliced genes correlated with AUD in our analyses were also linked with brain splicing associations with autism spectrum disorder and schizophrenia^2^, which included glutamate receptor (*GRIA2*) and calcium signaling genes (*CACNA1G, CAMK2D* and *CAMKMT*) as well as intracellular processes (*AKAP13, ARPP21, PRKACB* and *PTPRS*) and synaptic plasticity genes (*ARHGEF10L, ARHGEF4, CLASP2, GAPVD1, NTNG2, SUN1* and *TPM3*).

While our study characterizes the genetic roots of alcohol-related alternative mRNA splicing, we cannot dismiss the potential for alcohol-induced differential splicing. We found that many of the differentially spliced genes associated with AUD were also differentially spliced in primate models of chronic binge drinking. Notably, only five of these overlapping genes from analogous brain regions had a sQTL (2.9% of sQTLs). This may suggest both genetic and alcohol-related mechanisms underlying alternative mRNA splicing in the brain.

The current study should be interpreted in the context of the following limitations. Genetic effects from our study should be interpreted with caution and were limited to biallelic common SNPs. Polygenic scores used PRScise2, which chooses a threshold that maximizes and may over-fit the data. Additionally, polygenic prediction of splicing was done on the first principal component that explained < 5% of the variance in differential mRNA splicing for each brain region. Partitioned heritability analyses indicated that alternative mRNA splicing explained a significant amount of the heritability, but this is still ~2% of the total individual differences in problematic alcohol use and may include non-splicing related DNA variants. We are reticent to interpret individual sQTLs as these analyses were based on small samples. But, we utilized multiple tissue types and cross-referenced findings with GWASs with much larger samples. The GWASs used in our study included some overlapping participants (e.g., UK BioBank and Million Veterans Project) and were limited to individuals of European Ancestry.

In conclusion, we found a genetic component to brain-related alternative mRNA splicing underlying AUD and problematic alcohol use. We unveiled a host of genes that were differentially spliced between individuals with AUD and controls, which demonstrated stronger effects than classic differential expression analyses. By marshaling extant data sources with state-of-the-art methodology, we were able to make novel biological discoveries that added context to our genetic and neurobiological understanding of alcoholism.

## Acknowledgements

We acknowledge the National Institute on Drug Abuse award DP1DA042103 (awarded to RHCP). We are grateful for all of the participants from the GWAS studies as well as the public availability of the RNA-sequencing data. Without open sourced data or tools none of this research would have been possible.

## References

1. Pan Q, Shai O, Lee LJ, Frey BJ, Blencowe BJ. Deep surveying of alternative splicing complexity in the human transcriptome by high-throughput sequencing. Nat Genet 2008; 40(12): 1413–1415.

2. Gandal MJ, Zhang P, Hadjimichael E, Walker RL, Chen C, Liu S et al. Transcriptome-wide isoform-level dysregulation in ASD, schizophrenia, and bipolar disorder. Science 2018; 362(6420).

3. Donadoni M, Cicalese S, Sarkar DK, Chang SL, Sariyer IK. Alcohol exposure alters pre-mRNA splicing of antiapoptotic Mcl-1L isoform and induces apoptosis in neural progenitors and immature neurons. Cell Death Dis 2019; 10(6): 447.

4. Signor S, Nuzhdin S. Dynamic changes in gene expression and alternative splicing mediate the response to acute alcohol exposure in Drosophila melanogaster. Heredity (Edinb) 2018; 121(4): 342–360.

5. Farris SP, Arasappan D, Hunicke-Smith S, Harris RA, Mayfield RD. Transcriptome organization for chronic alcohol abuse in human brain. Mol Psychiatry 2015; 20(11): 1438–1447.

6. Lee C, Mayfield RD, Harris RA. Altered gamma-aminobutyric acid type B receptor subunit 1 splicing in alcoholics. Biol Psychiatry 2014; 75(10): 765–773.

7. Van Booven D, Mengying L, Sunil Rao J, Blokhin IO, Dayne Mayfield R, Barbier E et al. Alcohol use disorder causes global changes in splicing in the human brain. Transl Psychiatry 2021; 11(1): 2.

8. Zhou H, Sealock JM, Sanchez-Roige S, Clarke TK, Levey DF, Cheng Z et al. Genome-wide meta-analysis of problematic alcohol use in 435,563 individuals yields insights into biology and relationships with other traits. Nat Neurosci 2020; 23(7): 809–818.

9. Kranzler HR, Zhou H, Kember RL, Vickers Smith R, Justice AC, Damrauer S et al. Genome-wide association study of alcohol consumption and use disorder in 274,424 individuals from multiple populations. Nat Commun 2019; 10(1): 1499.

10. Palmer RH, McGeary JE, Heath AC, Keller MC, Brick LA, Knopik VS. Shared additive genetic influences on DSM-IV criteria for alcohol dependence in subjects of European ancestry. Addiction 2015; 110(12): 1922–1931.

11. Rao X, Thapa KS, Chen AB, Lin H, Gao H, Reiter JL et al. Allele-specific expression and high-throughput reporter assay reveal functional genetic variants associated with alcohol use disorders. Mol Psychiatry 2021; 26(4): 1142–1151.

12. Grant KA, Leng X, Green HL, Szeliga KT, Rogers LS, Gonzales SW. Drinking typography established by scheduled induction predicts chronic heavy drinking in a monkey model of ethanol self-administration. Alcohol Clin Exp Res 2008; 32(10): 1824–1838.

13. Baker EJ, Farro J, Gonzales S, Helms C, Grant KA. Chronic alcohol self-administration in monkeys shows long-term quantity/frequency categorical stability. Alcohol Clin Exp Res 2014; 38(11): 2835–2843.

14. Bolger AM, Lohse M, Usadel B. Trimmomatic: a flexible trimmer for Illumina sequence data. Bioinformatics 2014; 30(15): 2114–2120.

15. Dobin A, Davis CA, Schlesinger F, Drenkow J, Zaleski C, Jha S et al. STAR: ultrafast universal RNA-seq aligner. Bioinformatics 2013; 29(1): 15–21.

16. Browning BL, Zhou Y, Browning SR. A One-Penny Imputed Genome from Next-Generation Reference Panels. Am J Hum Genet 2018; 103(3): 338–348.

17. Choi SW, Mak TS-H, O’Reilly PF. Tutorial: a guide to performing polygenic risk score analyses. Nature Protocols 2020; 15(9): 2759–2772.

18. Li YI, Knowles DA, Humphrey J, Barbeira AN, Dickinson SP, Im HK et al. Annotation-free quantification of RNA splicing using LeafCutter. Nat Genet 2018; 50(1): 151–158.

19. Smedley D, Haider S, Ballester B, Holland R, London D, Thorisson G et al. BioMart--biological queries made easy. BMC Genomics 2009; 10: 22.

20. Love MI, Huber W, Anders S. Moderated estimation of fold change and dispersion for RNA-seq data with DESeq2. Genome Biol 2014; 15(12): 550.

21. Choi SW, O’Reilly PF. PRSice-2: Polygenic Risk Score software for biobank-scale data. Gigascience 2019; 8(7).

22. Wang K, Li M, Hakonarson H. ANNOVAR: functional annotation of genetic variants from high-throughput sequencing data. Nucleic Acids Res 2010; 38(16): e164.

23. Zhao H, Sun Z, Wang J, Huang H, Kocher JP, Wang L. CrossMap: a versatile tool for coordinate conversion between genome assemblies. Bioinformatics 2014; 30(7): 1006–1007.

24. Li YI, Wong G, Humphrey J, Raj T. Prioritizing Parkinson’s disease genes using population-scale transcriptomic data. Nat Commun 2019; 10(1): 994.

25. Barbeira AN, Pividori M, Zheng J, Wheeler HE, Nicolae DL, Im HK. Integrating predicted transcriptome from multiple tissues improves association detection. PLoS Genet 2019; 15(1): e1007889.

26. Barbeira AN, Dickinson SP, Bonazzola R, Zheng J, Wheeler HE, Torres JM et al. Exploring the phenotypic consequences of tissue specific gene expression variation inferred from GWAS summary statistics. Nat Commun 2018; 9(1): 1825.

27. Liu M, Jiang Y, Wedow R, Li Y, Brazel DM, Chen F et al. Association studies of up to 1.2 million individuals yield new insights into the genetic etiology of tobacco and alcohol use. Nat Genet 2019; 51(2): 237–244.

28. Zhou H, Rentsch CT, Cheng Z, Kember RL, Nunez YZ, Sherva RM et al. Association of OPRM1 Functional Coding Variant With Opioid Use Disorder: A Genome-Wide Association Study. JAMA Psychiatry 2020; 77(10): 1072–1080.

29. Johnson EC, Demontis D, Thorgeirsson TE, Walters RK, Polimanti R, Hatoum AS et al. A large-scale genome-wide association study meta-analysis of cannabis use disorder. Lancet Psychiatry 2020; 7(12): 1032–1045.

30. Schlesinger F, Tammena D, Krampfl K, Bufler J. Desensitization and resensitization are independently regulated in human recombinant GluR subunit coassemblies. Synapse 2005; 55(3): 176–182.

31. Penn AC, Balik A, Wozny C, Cais O, Greger IH. Activity-mediated AMPA receptor remodeling, driven by alternative splicing in the ligand-binding domain. Neuron 2012; 76(3): 503–510.

32. Acosta G, Freidman DP, Grant KA, Hemby SE. Alternative splicing of AMPA subunits in prefrontal cortical fields of cynomolgus monkeys following chronic ethanol self-administration. Front Psychiatry 2011; 2: 72.

33. Heck J, Parutto P, Ciuraszkiewicz A, Bikbaev A, Freund R, Mitlöhner J et al. Transient Confinement of Ca(V)2.1 Ca(2+)-Channel Splice Variants Shapes Synaptic Short-Term Plasticity. Neuron 2019; 103(1): 66–79.e12.

34. Chen L, Tian L, MacDonald SH, McClafferty H, Hammond MS, Huibant JM et al. Functionally diverse complement of large conductance calcium- and voltage-activated potassium channel (BK) alpha-subunits generated from a single site of splicing. J Biol Chem 2005; 280(39): 33599–33609.

35. Raj T, Li YI, Wong G, Humphrey J, Wang M, Ramdhani S et al. Integrative transcriptome analyses of the aging brain implicate altered splicing in Alzheimer’s disease susceptibility. Nat Genet 2018; 50(11): 1584–1592.

36. Takata A, Matsumoto N, Kato T. Genome-wide identification of splicing QTLs in the human brain and their enrichment among schizophrenia-associated loci. Nat Commun 2017; 8:14519.

37. Zhang X, Joehanes R, Chen BH, Huan T, Ying S, Munson PJ et al. Identification of common genetic variants controlling transcript isoform variation in human whole blood. Nat Genet 2015; 47(4): 345–352.

38. Pei W, Huang Z, Wang C, Han Y, Park JS, Niu L. Flip and flop: a molecular determinant for AMPA receptor channel opening. Biochemistry 2009; 48(17): 3767–3777.

39. Gilpin NW, Herman MA, Roberto M. The central amygdala as an integrative hub for anxiety and alcohol use disorders. Biol Psychiatry 2015; 77(10): 859–869.

40. Janak PH, Tye KM. From circuits to behaviour in the amygdala. Nature 2015; 517(7534): 284–292.

