## Supplementary Information for "Genome- and Transcriptome-wide Splicing Associations with Problematic Alcohol Use and Alcohol Use Disorder"

**Supplementary Figure S1** Principal Components Plot Showing Genetic Ancestry of Post-mortem RNA-sequencing data

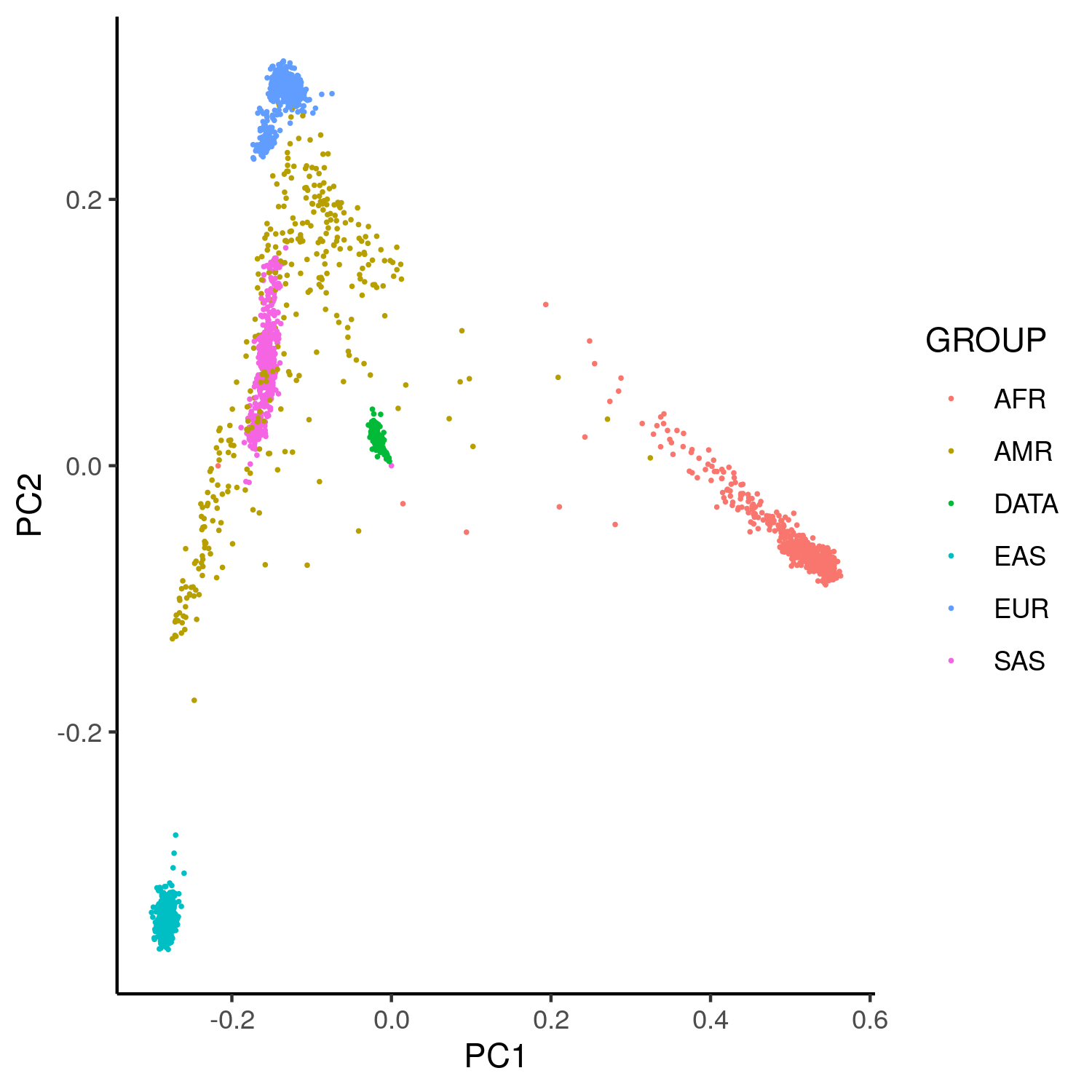

Each dot represents an individual sample and is color-coded by ancestry. Note that our RNA-seq data are shown in green, whereas all other colors represent ancestries from reference samples in 1K Genomes. AFR = African; AMR = Ad mixed American; EAS = Eastern Asian; EUR = European; SAS = South Asian.

**Supplementary Figure S2** Polygenic prediction of AUD and Differential Splicing with Clumping

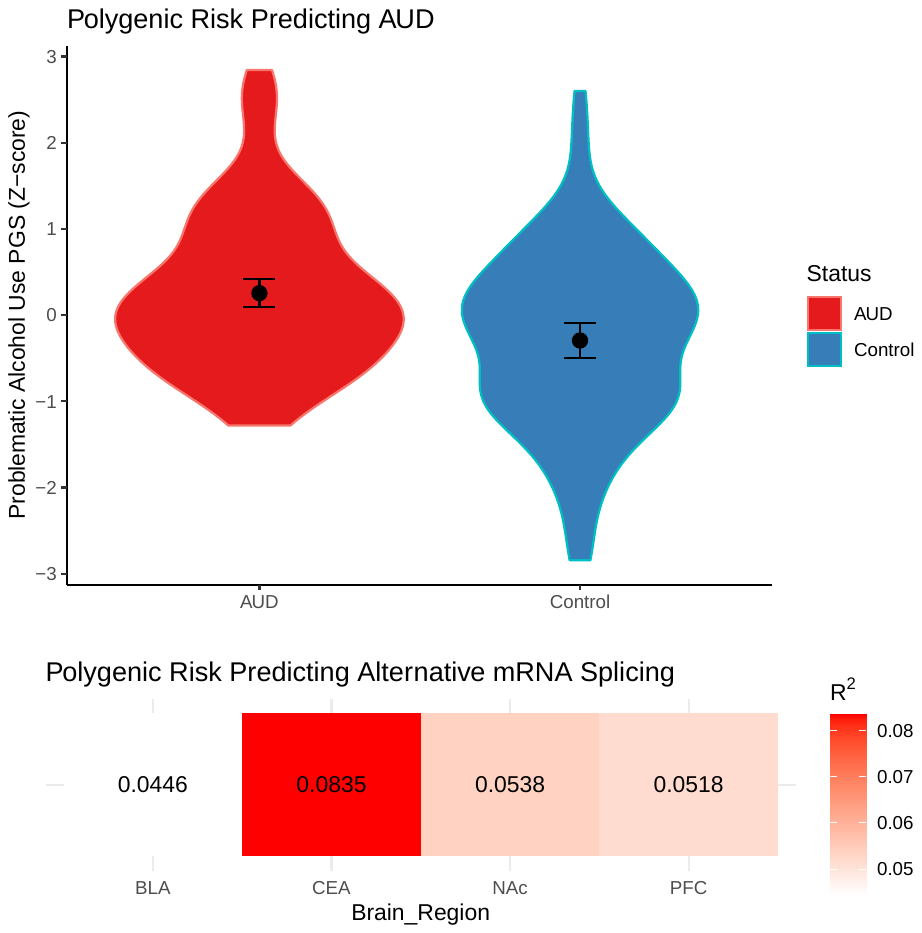

**A)** Violin plot showing polygenic score distributions of problematic alcohol use between individuals with AUD and controls. Mean and standard error are shown. **B)** Heat matrix showing the amount of variance explained (R^2^) by polygenic prediction of differential splicing results for each brain region. Principal components (PC) analysis was used to distil differential splicing results into a single metric (1^st^ PC).

**Supplementary Figure S3** Differentially Spliced Addiction Genes in the CEA

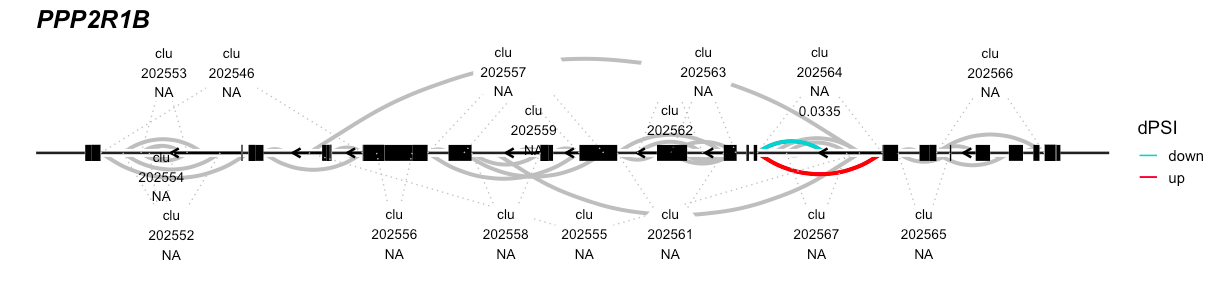

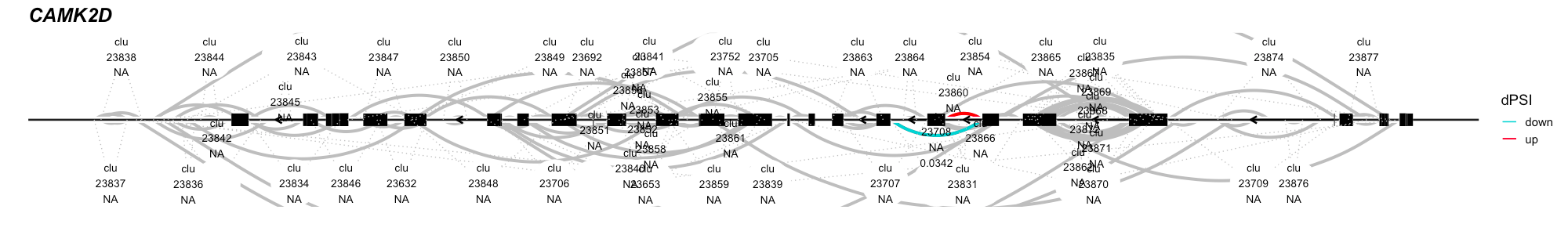

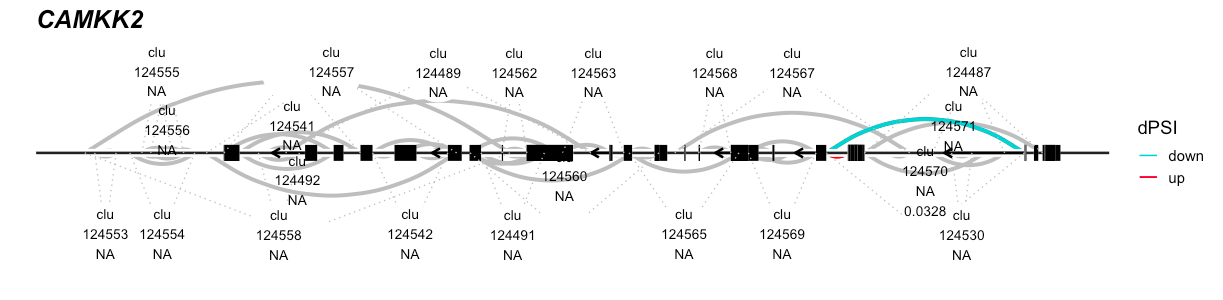

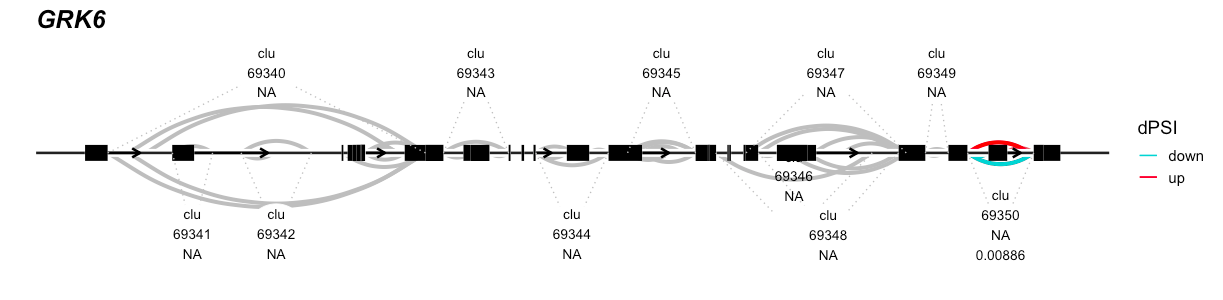
Differentially spliced clusters in the genes are labeled and emphasized in color. Red arcs show increased splicing in individuals with AUD (e.g., gene exons are more likely to be connected in AUD than controls). Cyan arcs represent decreased splicing in individuals with AUD (e.g., gene exons are more likely to be connected in controls than in individuals with AUD).

**Supplementary Figure S4** Differentially Spliced Addiction Genes in the BLA

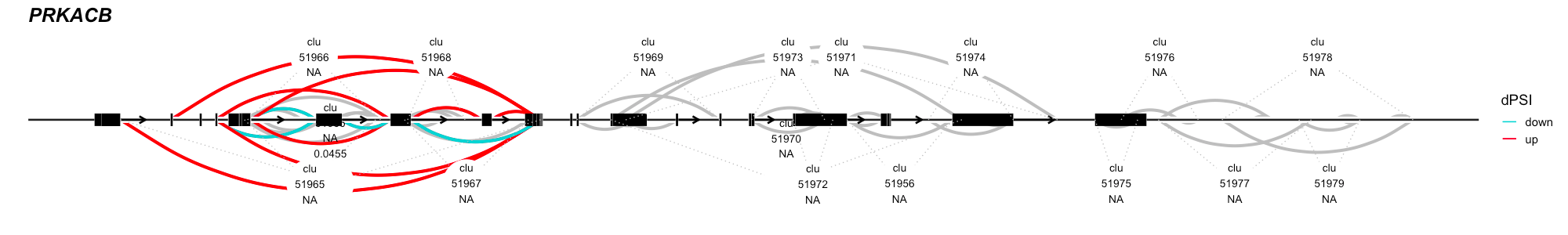

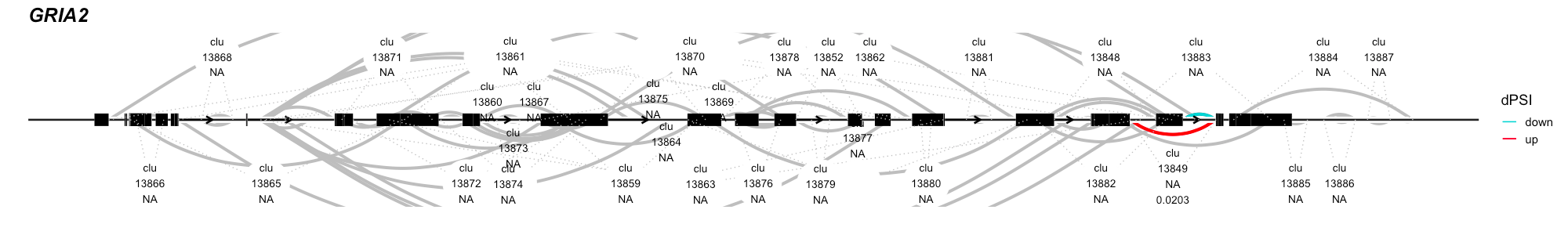

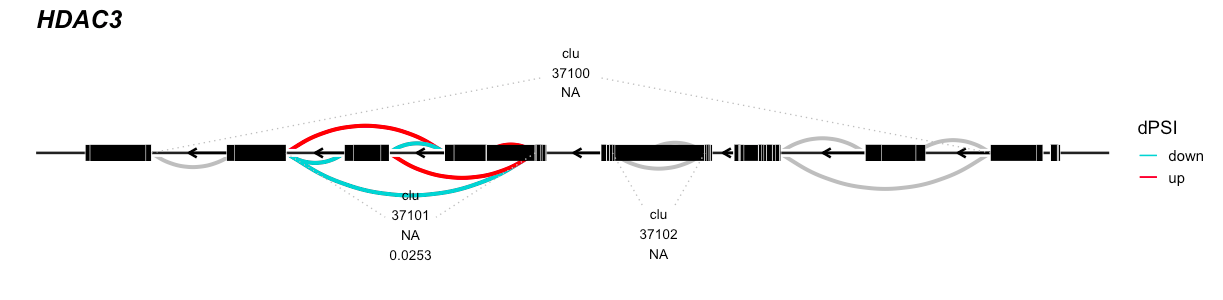

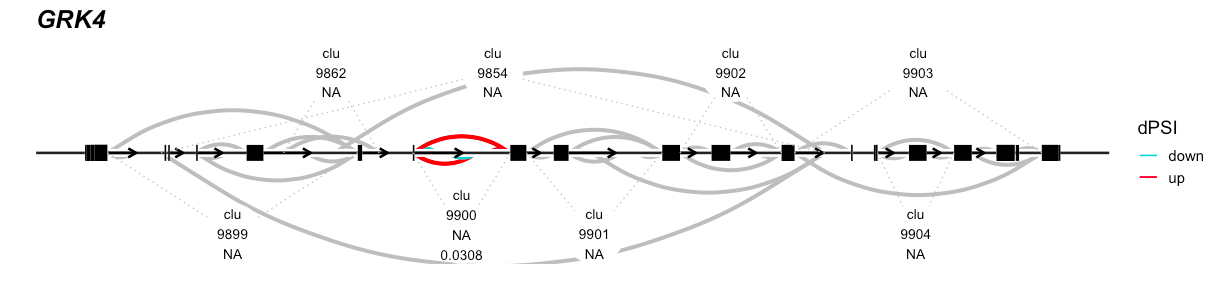
Differentially spliced clusters in the genes are labeled and emphasized in color. Red arcs show increased splicing in individuals with AUD (e.g., gene exons are more likely to be connected in AUD than controls). Cyan arcs represent decreased splicing in individuals with AUD (e.g., gene exons are more likely to be connected in controls than in individuals with AUD).

**Supplementary Figure S5** Differentially Splicing of *GRIA2* in BLA Associated with AUD

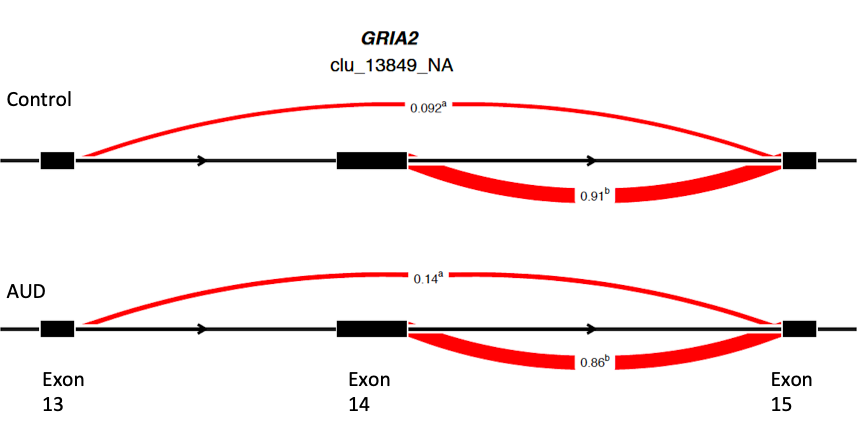

Exons are represented by black boxes and introns by black lines. Red arcs delineate splicing events and the numbers within each arc represent how often the exons are spliced together in either a control (top) or an AUD brain (bottom). Note that individuals with AUD are more likely to skip exon 14 in the BLA, which corresponds to an annotated splice site that influences AMPA receptor opening.

**Supplementary Figure S6** Differentially Spliced Genes Across Human AUD and Chronic Alcohol Use in Monkeys

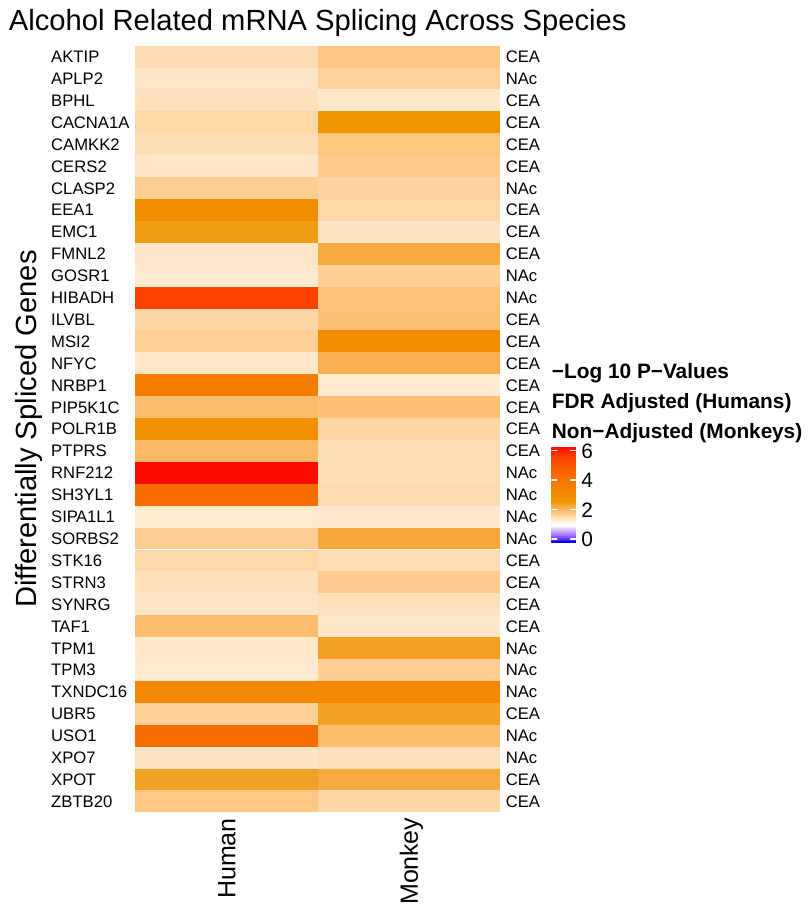

Heatmatrix plot showing differentially spliced genes associated with human AUD and monkey alcohol consumption. Note differential splicing in monkeys used a nominal p-value cutoff of p < 0.05 due to low power. Also, note that no overlap was found in the PFC across species and that the monkey data did not contain BLA data.

**Supplementary Figure S7** Splicing TWAS Associations across Substance Use Traits

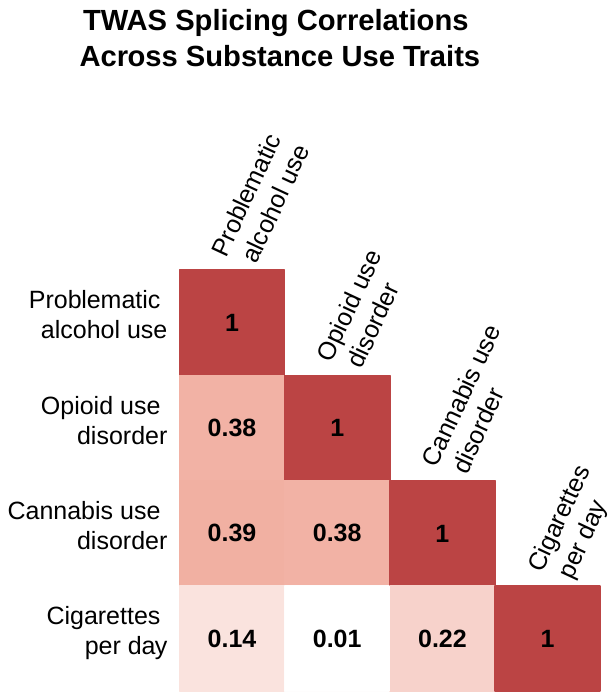

Heatmatrix plot showing the results of Correlations from the 1,397 significant Splicing TWAS associations (FDR < 0.05) across all substance use traits. Note only the correlation between cigarettes per day and opioid use disorder was non significant (p > 0.05).
